# LonP1 chaperone-like activity is ATPase independent and is mediated by its N-domain

**DOI:** 10.64898/2026.05.06.723147

**Authors:** Mette Ahrensback Roesgaard, Philipp Armbruster, Timothy Sharpe, Niko Schenck, Jan Pieter Abrahams

## Abstract

The mitochondrial Lon protease is essential for proteostasis through ATP-dependent proteolysis and suppression of protein aggregation through an unknown mechanism. Here we show in three independent aggregation systems that human Lon protease (LonP1) directly interacts with fibrillar aggregates to prevent further aggregation: LonP1 binds amyloid fibrils and inhibits their growth, independently of its protease and ATPase activities. This aggregation inhibition depends on hexamer stability, and even the N-domain hexamer of LonP1 lacking all catalytic domains inhibited aggregation, which localizes its fibril-binding interface. We propose that chaperone deficiencies in LonP1 mutants that are associated with genetic disease, are caused by reduced hexamer stability or increased turnover. Our results clarify the observed dual protease and chaperone function of LonP1 by localizing them to different domains and separating the catalytic activities, thereby facilitating targeting the specific functionalities. Further, we identify the structure of the chaperone substrate to be fibrillar aggregates, suggesting that LonP1 may protect against amyloid fibrils in healthy individuals.

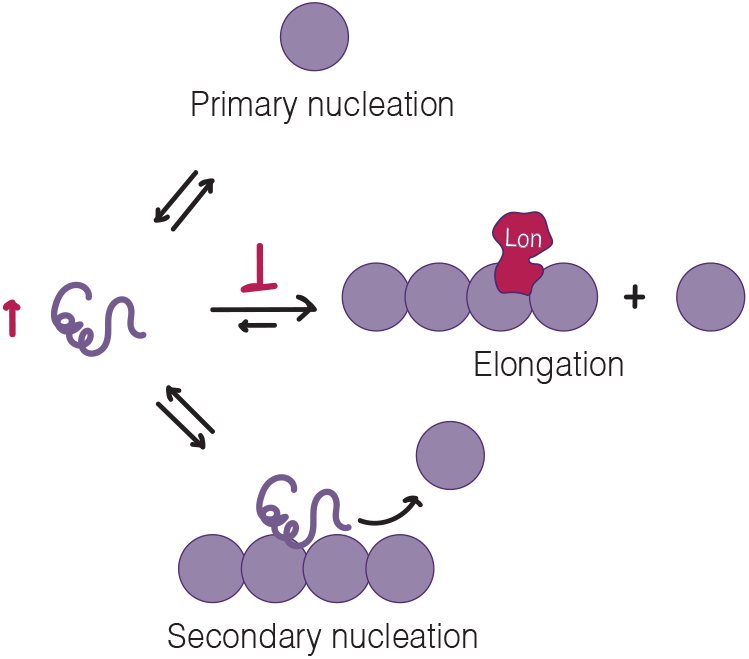

**Significance:** The mitochondrial Lon protease has long been proposed to function both as a protease and as a chaperone, though the mechanism of its chaperone activity is debated. Here, we show that human Lon binds to fibrillar protein aggregates and inhibits their elongation, but do not find evidence for chaperoning unfolded chains. Further, our findings challenge the current view that ATPase activity is required for Lon chaperone function. Instead, our results suggest that chaperone deficiency of Lon variants can be explained by variant stability. Our results provide a mechanistic separation of the protease and chaperone-like function LonP1, thereby opening up for targeting one of the functions specifically, and provide new insight into how Lon dysfunction may contribute in multiple ways to age-related and proteostasis-related diseases.

## Introduction

The Lon protease is a mitochondrial protease and ATPase that is essential for mitochondrial homeostasis (1-4). The Lon protease has several other functionalities including DNA binding (5), implications in mitochondrial import (6) and chaperoning (6-8). Disturbed mitochondrial homeostasis is a hallmark in neurodegenerative diseases (9). LonP1 inhibition increases pathology in Parkinson’s disease (10), and LonP1 expression levels are decreased in Alzheimer’s disease, with some evidence that LonP1 interacts directly with the amyloidogenic proteins (11). While the mechanism for the protease function of the human Lon protease (LonP1) is well described (12, 13), the mechanisms for its other functions remain unclear.

Evidence for Lon chaperone function was first observed in yeast (8) and later in mammalian cells, where knock-out and knock-down of LonP1 cause mitochondrial proteins to aggregate in the mitochondrial matrix (2, 4, 6, 7). Overexpression of proteolytically inactive Lon (S855A) rescues the appearance of mitochondrial protein aggregates, showing that Lon protease activity is not required for aggregation rescue, whereas Lon ATPase activity is required (6-8). Two known LonP1 disease variants, R301W and R721G, were found to be proteolytically active but chaperone inactive, and the latter variant also had decreased ATPase activity (14, 15).

However, these observations do not prove that LonP1 acts as a chaperone through direct interaction with unfolded proteins or aggregates. Since the Lon protease has several other functions, altering its activity disturbs multiple pathways that may influence aggregate formation. For example, knock-down of LonP1 causes higher transcription of mitochondrial proteins, likely caused by higher levels of the transcription factor TFAM, a proteolytic substrate of LonP1 (16, 17). Further, cells can compensate LonP1 knockdown by altering levels of other protein quality control components, such as mitochondrial import proteins and chaperones (18). Lastly, since proteolytically inactive LonP1 hydrolyses ATP at near wildtype activity (19), an overexpression could lead to a depletion of ATP in the mitochondria, influencing several homeostasis pathways. Together, these variables make it difficult to study LonP1 chaperone activity from cell culture experiments alone and highlights the need for *in vitro* approaches where variables are known and controllable.

Currently, LonP1 is proposed to chaperone unfolded substrates during import to the mitochondrial matrix (6), and nascent polypeptides during mitochondrial translation (7), though the molecular mechanism has not been demonstrated. In this work, we use chaperone assays, where the interaction between LonP1 and an unfolded protein substrate is probed. We find that the human Lon protease inhibits amyloid fibril growth *in vitro*, not by interacting with unfolded monomers, but by interacting with the fibril species. We discuss the chaperone function of LonP1 in light of these findings and propose that fibril elongation inhibition is the mechanism by which LonP1 prevents protein aggregation in the mitochondrial matrix.

Further, we consider the implications of this activity for the link between mitochondrial homeostasis and amyloid disease.

## Results

### LonP1 reduces aggregation in an in vitro expression assay by reducing total yield

We first tested a published *in vitro* assay for LonP1 chaperone activity, where an aggregation prone membrane protein (OXA1L) is expressed in the presence of proteolytically inactive LonP1 S855A (7). We used the same commercial kit and monitored the aggregation over time using Dynamic Light Scattering. We found that the average hydrodynamic diameter increased in the control reactions with Bovine Serum Albumin (BSA) (figure 1.A), with particles of a diameter of 100-300 nm appearing, consistent with aggregation (supplementary figure S1). In reactions with 2 µM proteolytically inactive LonP1 S855A, the aggregation is completely surpressed (figure 1A, supplementary figure S1). However, we found that the total yield of the expression was 4-fold decreased in the presence of 2 µM LonP1, even when expressing the soluble protein DHFR, and non-detectable for expression of OXA1L (figure 1.B). The ATPase deficient variant of LonP1 (K529R) did not decrease the yield (figure 1.D), and rescued aggregation as measured by DLS, though not to the same extent as LonP1 S855A (figure 1A). Adding an inhibitor of LonP1 ATPase activity, CDDO-Me (20), doubled the yield for expressions with ATPase active LonP1 (figure 1.C). Thus, the ATPase activity of LonP1strongly inhibits the expression assay, and the effect of the ATPase activity and the chaperone-like activity can therefore not be separated in this assay.

**Figure 1:**
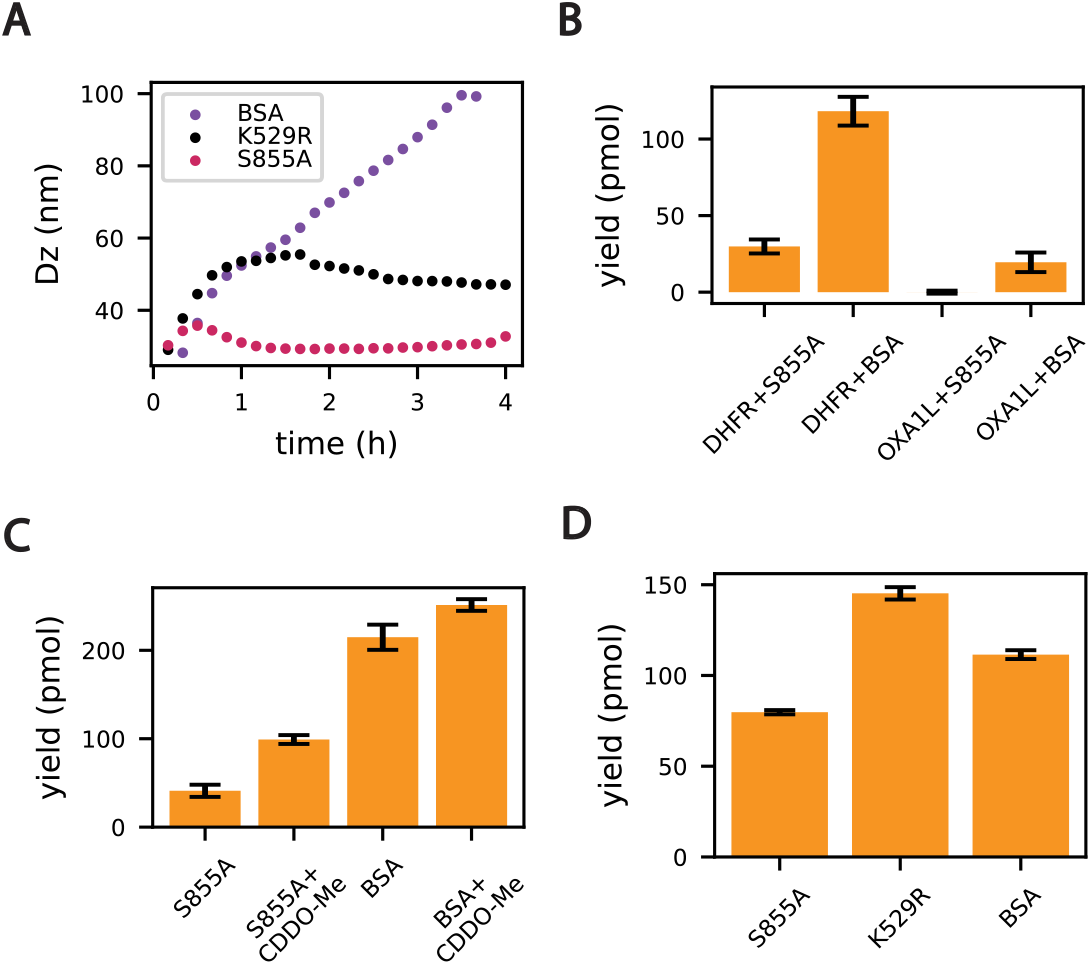
**A**. Intensity-weighted average hydrodynamic diameter (D_z_) over time of an *in vitro* expression of OXA1L in the presence of 2 µM LonP1 S855A, LonP1 K529R or BSA, measured with DLS. **B**. Total yield of *in vitro* expression of OXA1L or soluble DHFR in the presence of 2 µM LonP1 S855A or BSA. **C**. Total yield of *in vitro* expression of DHFR in the presence of 1 µM LonP1 S855A or 2 µM BSA, with and without 5 µM CDDO-Me. **D**. Total yield of *in vitro* expression of DHFR in the presence of 0.7 µM LonP1 S855A, LonP1 K529R or 2 µM BSA. Error bars on B-D represenst standard error of the mean of duplicate measurements.

### Proteolytically inactive LonP1 inhibits amyloid-beta fibril elongation and distorts fibril morphology

Amyloid-beta 42 is a natively unfolded protein fragment that undergoes spontaneuos aggregation into amyloid fibrils at neutral pH, and can therefore be used as a simple system to study aggregation inhibition. Using the amyloid dye Thioflavin T (ThT), we monitored amyloid-beta 42 in the presence of increasing concentrations of proteolytically inactive LonP1 S855A or BSA as a control, and found that LonP1 S855A causes a concentration-dependent delay in fibrillization (figure 2.A). Kinetic analysis using Amylofit (21) showed that the best fit was obtained when using an elongation inhibition model. Rate constants for primary nucleation, fibril elongation and secondary nucleation were determined from reactions with only amyloid-beta and fixed while fitting inhibition constants from reactions with LonP1 present (figure 2.B). Elongation inhibiton dominated the fit with an inhibition constant in the nanomolar range (5e-10), and resulted in the best fit, though a combination of secondary and primary nucleation inhibition explains the data equally well. No delay in the onset of fibrillization was observed, and the fit for primary nucleation inhibition was poor, indicating that LonP1 does not interact with the unfolded monomer.

**Figure 2:**
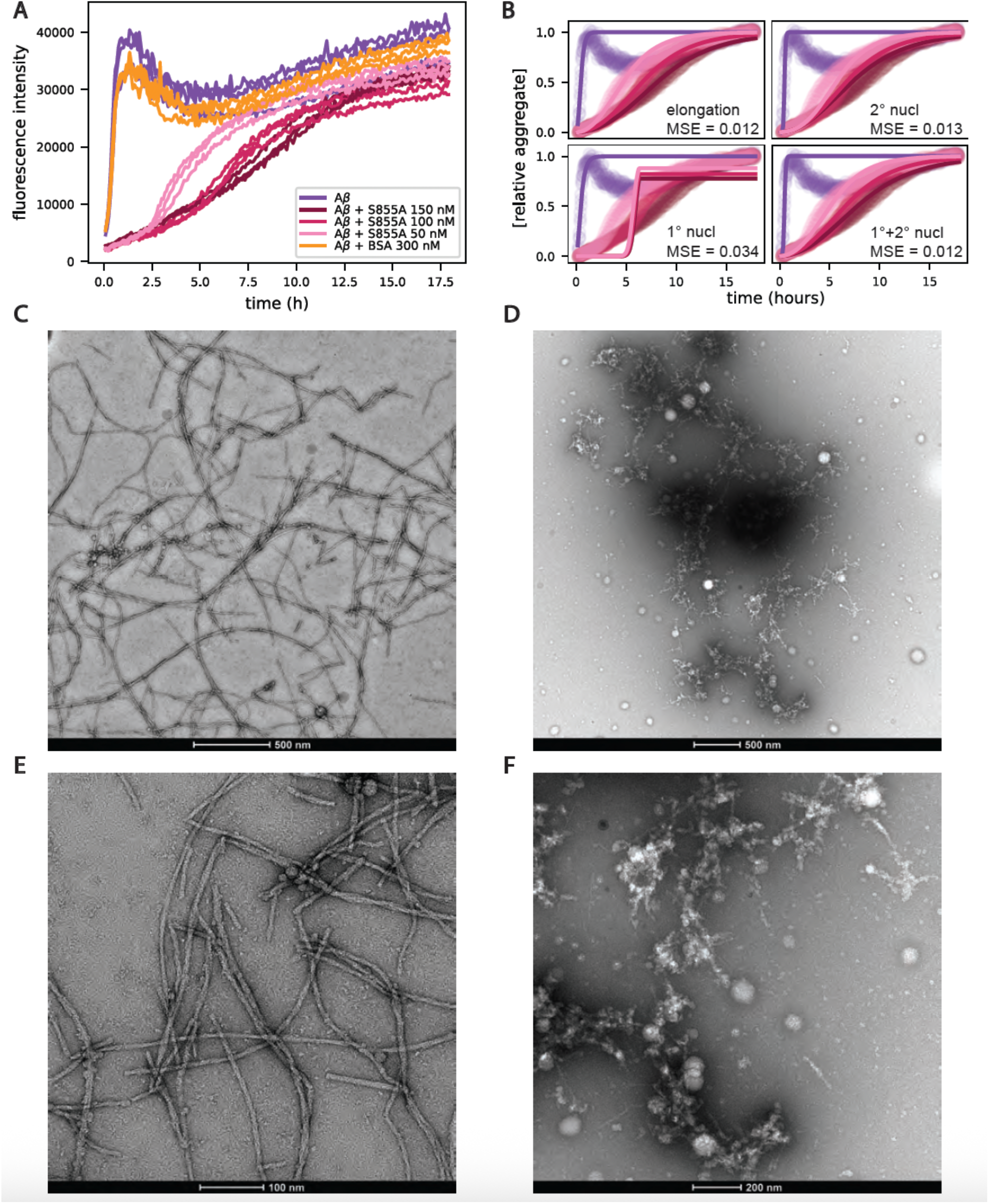
**A**. Fibrilization of amyloid-beta over time measured by Thioflavin T fluorescence, alone and in the presence of 300 nM BSA (control) and 50 nM, 100 nM or 150 nM LonP1 S855A (hexamer concentration), absolute intensity with background fluorescence subtracted, triplicate reactions. **B**. Fits to the data in A with the inhibition constant of elongation, primary nucleation, secondary nucleation or both secondary and primary nucleation as free parameters and mean square error (MSE) reported on each fit. **C-F**: Negative stain electron microscopy images of amyloid-beta fibrils after 18 hours of incubation in the presence of either 300 nM BSA as a control (C,E) or 50 nM LonP1 S855A (D,F).

At reaction endpoints, samples were analyzed by negative stain electron microscopy. Both reactions with LonP1 S855A and with the control, BSA, had fibrillar structures. However, the control reaction has well defined, twisted and mature fibrils, while amyloid fibrils formed in the presence of LonP1 are thinner, shorter and proto-filament-like (figure 2 C-F), indicating that LonP1 inhibits elongation of amyloid-beta fibrils and distorts their morphology. Fibrils formed in the presence of LonP1 are decorated with particles similar in size to free LonP1 observed on the grid, suggesting that LonP1 binds to the fibril surface. However, these particles could also be secondary nucleation events, and the limited resolution of negative-stain electron microscopy does not allow us to discriminate with certainty.

### LonP1 inhibits insulin fibril elongation and associates with fibril species

Reduction of insulin with DTT dissociates the A and B chain, causing rapid and effectively irreversible aggregation of the unstable B chain. We find that LonP1 S855A inhibits insulin aggregation, not by delaying the onset, but by lowering the plateau of the aggregation profiles as measured by absorbance (figure 3.A), meaning that LonP1 must either reduce the fibril mass or change the fibril size. Total fibril mass, measured by ThT, did not change in the presence of LonP1 (figure 3.B), whereas DLS reveals that most fibrils grow slower and are shorter if they were formed in the presence of LonP1 (figure 3.C). Together, these results show that LonP1 presence increases the number of insulin fibrils and reduces their length. For reduced insulin fibrillization kinetics, elongation inhibition is expected to increase nucleation, because the unfolded state is not stable. These results therefore further support an elongation inhibiton function of LonP1. Additionally, LonP1 co-precipitates with both insulin fibrils and amyloid-beta fibrils, indicating association with the fibril species (supplementary figure S2).

**Figure 3:**
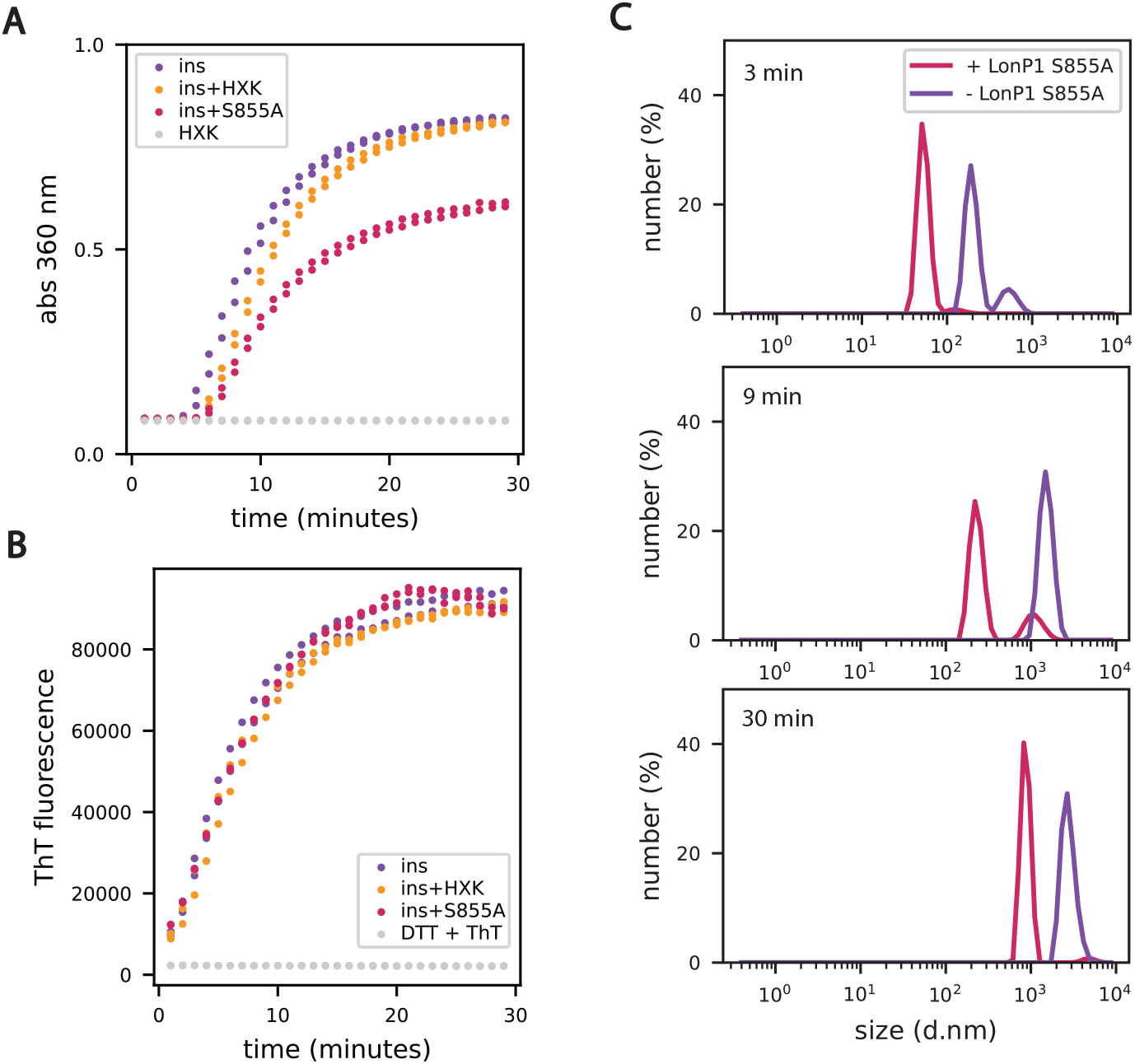
**A-B**. Insulin aggregation over time induced by DTT in the presence of 150 nM LonP1 S855A or 1 µM hexokinase (HXK), measured by absorbance at 360 nm (A) or Thioflavin T fluorescence (B), duplicate reactions. **C**. Size distribution time points of aggregating insulin with or without 100 nM LonP1 S855A, measured with DLS.

### Inhibition of fibril elongation depends on LonP1 stability, not on ATPase activity

We tested proteolytic inactive LonP1 (S855A), ATPase deficient LonP1 (K529R) and the disease variants R301W and R721G (14, 15) (supplementary figure S3) for their ability to inhibit insulin fibril growth. All variants decreased insulin aggregation at nanomolar concentrations when using hexameric LonP1 isolated with Size Exclution Chromotography. Addition of ATP did not affect aggregation inhibition of ATPase active LonP1 S855A, nor on ATPase inactive LonP1 K529R (figure 4.A). Because nucleotide binding is known to stabilise LonP1 (22), we tested if the aggregation inhibition depends on variant stability rather than on ATPase activity, by incubating hexameric LonP1 S855A and LonP1 K529R with or without ADP overnight at 37 °C prior to the aggregation assay. LonP1 S855A retained aggregation inhibition activity when incubated with ADP, but loses it in the absence of ADP, while LonP1 K529R inhibtion activity cannot be fully rescued by the presence of ADP (figure 4. B), indicating a correlation with variant stability rather than ATPase activity. Next, we found that the LonP1 N-domain alone can rescue insulin aggregation, but at much higher concentrations than what has been observed for the Lon bacterial homologue (23), and at about a 10-fold higher molar concentration than the full length variants of LonP1 (figure 4. C). This indicates that the fibril-LonP1 interaction occurs at the LonP1 N-domain. Under these conditions, only 10% of the N-domain is hexameric, as measured by mass photometry (supplementary figure S4) which further indicates that hexamer stability is required for aggregation inhibition.

**Figure 4.**
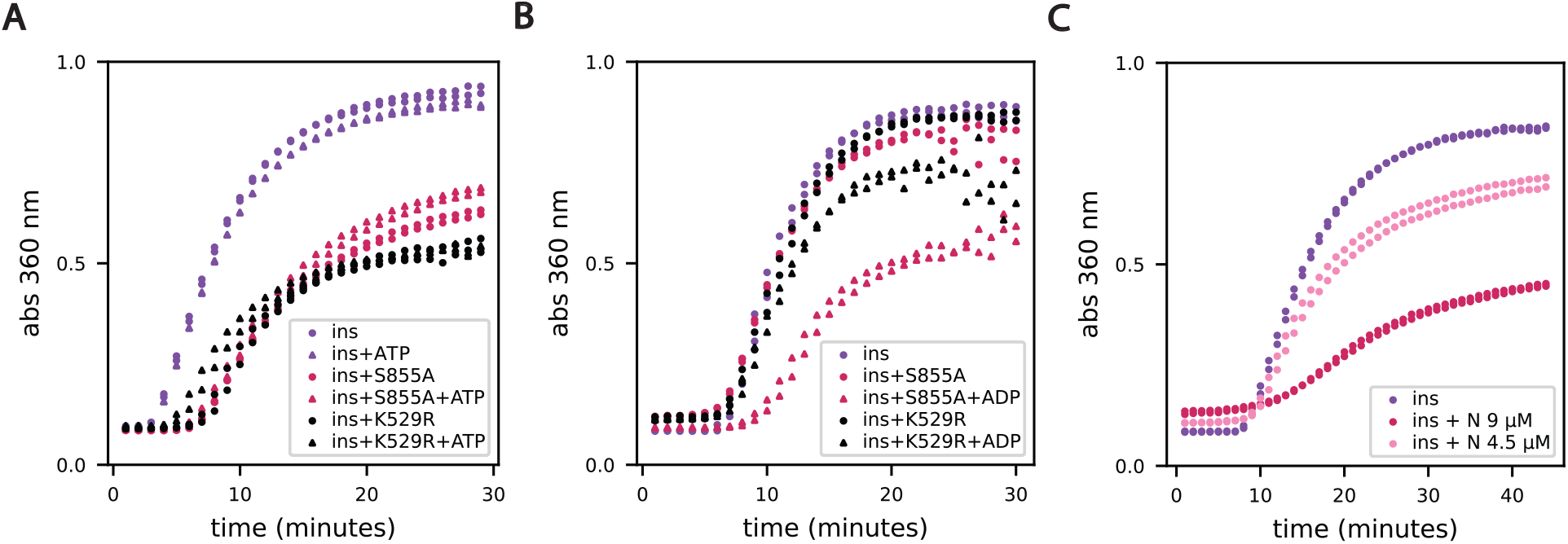
Insulin aggregation over time induced by DTT in the presence of LonP1 variants, measured by absorbance at 360 nm, duplicate reactions. **A.** Effect on insulin aggregation of 150 nM LonP1 S855A or LonP1 K529R, with or without 1 mM ATP. **B**. Insulin aggregation prevention of 150 nM LonP1 S855A and K529R after overnight incubation with or without 1 mM ADP at 37 °C. **C**. Insulin aggregation with LonP1 N-domain at 4.5 or 9 µM concentration.

## Discussion

We do not find evidence for chaperoning activity of the Lon protease on unfolded proteins in our kinetic analysis of aggregation curves, nor are we able to determine that the decrease in aggregation of a nascently transcribed protein chain in the presence of LonP1 is caused by chaperoning and not by inhibition of the *in vitro* expression system. We conclude that the ATPase activity of LonP1 strongly inhibits the expression assay, obscuring interpretations regarding ATP hydrolysis as a requirement for chaperoning, but note that some aggregation rescue can be observed for ATPase deficient LonP1.

Instead, our data supports a model where LonP1 rescues aggregation by direct interaction with fibril species, causing elongation inhibition. LonP1 exhibits this activity on two amyloidogenic proteins with different primary structure, which indicates that it may recognize the common secondary structure element, the amyloid cross beta fold(24), or the disordered regions of the fibrils, rather than a specific amino acid sequence.

Amyloid-beta fibrils have previously been observed to inhibit LonP1 proteolytic activity *in vitro* (11). In light of our results, this is likely due to LonP1 association with amyloid-beta fibrils and thereby removal from the pool of soluble, proteolytically active LonP1. Thus, LonP1 and amyloid-beta fibrils appear to exhibit reciprocal inhibition.

This raises the question whether the interaction between amyloids and Lon is functional or a disease symptom. It is well established that mitochondrial homeostasis is disturbed in amyloid diseases through multiple pathways, and there is some indications that mitochondrial dysfunction cause amyloid formation (25). Expression levels of Lon decrease during ageing (26), and are reduced in Alzheimer cells with amyloid plaques (27), again indicating that lack of Lon may contribute to amyloid formation. As a protease, Lon clears abundant and misfolded protein (28), and its reduced expression can be expected to cause more misfolded and aggregation prone proteins, but the results we show here suggest that Lon may additionally directly function as a protecter against amyloid growth. Interestingly, for Parkinson’s disease, an increase of LonP1 levels was found to correlate with the accumulation of alpha synuclein aggregates (29), although treating cells with the LonP1 inhibitor CDDO-Me resulted in an increase in alpha synuclein fibrils (10).

Contrary to previous studies (7, 14) we do not find that the disease variants LonP1 R301W and LonP1 R721G, and the ATPase deficient LonP1 K529R, are unable to rescue aggregation. Instead, we show that LonP1 hexameric stability is required for aggregation rescue, and that even the LonP1 N-domain that does not have and ATPase or a protease site, can rescue aggregation in its hexameric state. Consistent with this, the disease variant R721G was mostly not hexameric (14). This may explain why some variants fail to rescue aggregation in cell cultures, although being active *in vitro*.

Our results provide a mechanistic understanding for the distinguishment of substrates for proteolysis or chaperoning: the chaperone substrate is the amyloid fibril, which LonP1 attaches to through an interaction between the N-domain and the fibril surface. The substrate is unlikely to be unfolded due to the strong and inaccesible hydrogen bonding in the cross beta fold and would not be able to pass through the Lon catalytic chamber, where proteolysis takes place. As LonP1 is a cancer drug target (30), we believe that our results are of high interest for researchers targeting specifically its proteolytic or chaperone function. We strongly suggest caution in stating that a reduced ATPase activity of Lon equals a reduced chaperone function, and encourage further *in vivo* research to determine if the fibril elongation inhibition function and the chaperone-like function of LonP1 are indeed the same function.

## Methods

### Expression and purification of LonP1 wild type and variants

LonP1 wildtype and full length variants (R301W, K529R, R721G, S855A) were expressed and purified according to the published protocol from Mohammed et al (12), with the following changes; cultures were grown in pH-buffered Terrific Broth media, precultures were grown at 30 °C, expression was induced with 0.4 mM IPTG, lysozyme (Roche) was added during cell lysis and no EDTA or reducing agent was used in the purification. The N-domain of LonP1 (residues 115-409) was expressed and purified as the wild type, but with the addition of an ion exchange step, where the purified protein was buffer exchanged into 20 mM bis-tris propane pH 8, 100 mM NaCl and 5 mM MgCl2, and loaded onto a capto Q anion exchange column and eluted at 36% 1 M NaCl elution buffer. All plasmids were purchased from GeneScript.

### Dynamic Light Scattering of in vitro expression and yield determination

*In vitro* expression of the membrane protein OXA1L (72-435) and soluble protein DHFR was performed with the New England Biolabs PURExpress® In Vitro Protein Synthesis kit. The plasmid for DHFR was provided with kit, and the plasmid for OXA1L was purchased from GeneScript. The expression reaction was assembled according to the protocol of the kit, with the addition of either Bovine Serum Albumin (BSA) (Sigma-Aldrich, CAS 9048-46-8) or LonP1 to a total volume of 30 µl. The reaction was immediately transferred to a quartz cuvette (ZEN2112 3 mm, Malvern Panalytical) and Dynamic Light Scattering was measured in a Malvern Zetasizer Nano DLS, with 50 measurements of each 5 seconds per time point, measurement position 4.20 mm and attenuator 6. The reaction was run at 37 °C without shaking. The total yield was determined using TCA precipitaion and radio-labeled Methionine S-35 (from Hartmann Analytic, translation grade, specific activity 1000 Ci/mmol, radioactive concentration 740MBq (20mCi)/ml), according to the protocol from the PURExpress kit. CDDO-Me was aquired from a collaborator.

### Amyloid-beta aggregation inhibition assay

Synthetic Amyloid β Protein Fragment 1-42 from Sigma-Aldrich (CAS 107761-42-2) was dissolved in 50 mM NaOH, pH 11.5, sonicated, and aliqouted and stored at -80 °C until use.

To start the fibrilization, amyloid-beta aliqouts were diluted to 3 µM in a buffer containing 50 mM HEPES, 150 mM NaCl, 5 mM MgCl2, and 10 µM Thioflavin T (Calbiochem, CAS 2390-54-7) at pH 7.5, in the presence of 50-150 nM LonP1 S855A or 300 nM BSA (Sigma-Aldrich, CAS 9048-46-8), at a volume of 100 µl. Reactions took place at 37 degrees C with no shaking in a sealed 96-well half area clear bottom black non-binding plate (Corning 3881). Thioflavin T fluorescence was measured every 5 minutes for 18 hours with a 450 nm excitation filter and a 485 emission filter using using a Tecan Spark Multi-Mode Plate-Reader.

### Insulin aggregation inhibition assay

Following the method described by (31), bovine insulin (Sigma-Aldrich, CAS 11070-73-8) was diluted to a final concentration of 600 mg/ml (100 µM) into a buffer containing 50 mM HEPES, 150 mM NaCl, 5 mM MgCl2 at pH 7.5, and provoked to aggregate by adding 20 mM Dithiothreitol (DTT), in the presence of purified LonP1 variants at various concentration or Hexokinase from S. cerevisiae (H4502, Sigma-Aldrich). Aggregation was meausered in 96-well transparent plates by absorbance at 360 nm in a Tecan Spark Multi-Mode Plate-Reader every minute for 45-60 minutes, at a temperature of 25 degrees C, without shaking.

Insulin was added to the reactions with a multi-channel pipette as a last step, to start all reactions simultaneously.

### Dynamic Light Scattering of insulin aggregation

600 mg/ml bovine insulin (Sigma-Aldrich, CAS 11070-73-8), 20 mM DTT and 0 or 100 nM LonP1 S855A were mixed and 40 µl were transferred to a quartz cuvette (ZEN2112 3 mm, Malvern Panalytical). After 5 minutes incubation, Dynamic Light Scattering measurements were made every 3 minutes for 30 minutes on a Malvern Zetasizer Nano DLS at measurement position 4.20 mm and attenuator setting 6. Each timepoint is 20 measurements of 5 seconds.

### Co-precipitation

Samples at the end points of insulin or amyloid-beta reactions or samples of incubations with LonP1 or alpha-synuclein were centrifuged for 5 minutes at 13,000 × g. Pellet and supernatant were separated, and the pellet was redissolved in buffer to the same volume as the supernatant. Samples from the resuspended pellet and the supernatant were mixed with Laemmli buffer and loaded on NuPAGE 4-12% Bis-Tris Gels from ThermoFisher. Gels were stained with QuickBlue Protein Stain (Lubio science) and imaged on a Vilber Lourmat fusion FX imager.

### Negative stain electron microscopy of fibrils

During amyloid-beta Thioflavin T measurements, equivalent reactions without Thioflavin T at a volume of 100 µl were run simultaneuosly in test tubes at 37 degrees C without shaking, and a sample was taken for negative stain electron microscopy at 0 h, 2 h, 5 h, and 18 h.

Reactions were mixed gently by pipetting up and down before taking a sample to resuspend precipitated protein without destroying fibril morphology. 5 µl sample was deposited on carbon coated parlodion grids, incubated for 1 minute, blotted from the side, washed, and stained with uranyl-acetate. Images were aquired on a Thermo Fisher Scientific™ Talos L120C G2 electron microscope with a Thermo Fisher Scientific Ceta 16M CCD detector.

## Acknowledgement

We thank the BioEM Lab of the Biozentrum, University of Basel, and Carola Alampi for their support with the negative stain experiments, the Biophysics Facility at the Biozentrum, University of Basel, and our Marcelina Sikora and Natalie Peter for technical support.

## Author contributions

Roesgaard, M. A. performed the experiments, directed the project and wrote the manuscript. Sharpe, T. contributed to the design and execution of experiments. Armbruster, P. established a purification protocol for the N-domain. Schenck, N. manufactured materials and discussed the progress of the project. Abrahams, J. P. directed the project, provided funding and revised the manuscript. All authors have discussed the results and commented on the manuscript.

## Supplementary Material

**Supplementary figure S1:**
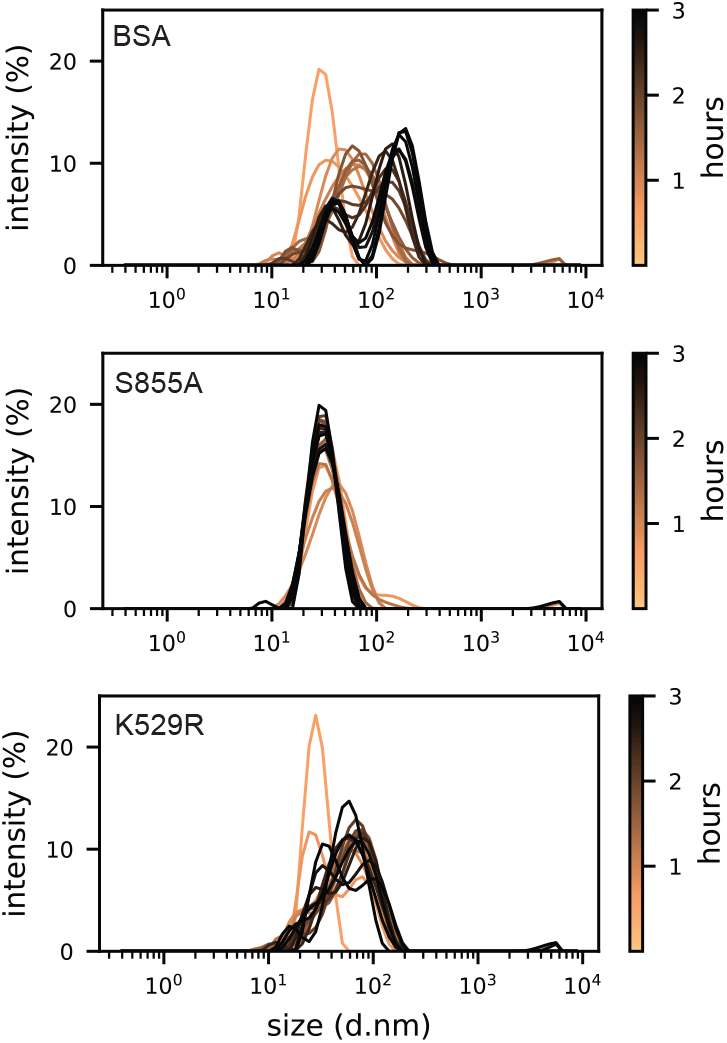
Size distribution over time in terms of intensity, which is sensitive to larger particles, of *in vitro* expression of OXA1L in the presence of 2 µM LonP1 S855A, LonP1 K529R or BSA, measured with Dynamic Light Scattering.

**Supplementary figure S2:**
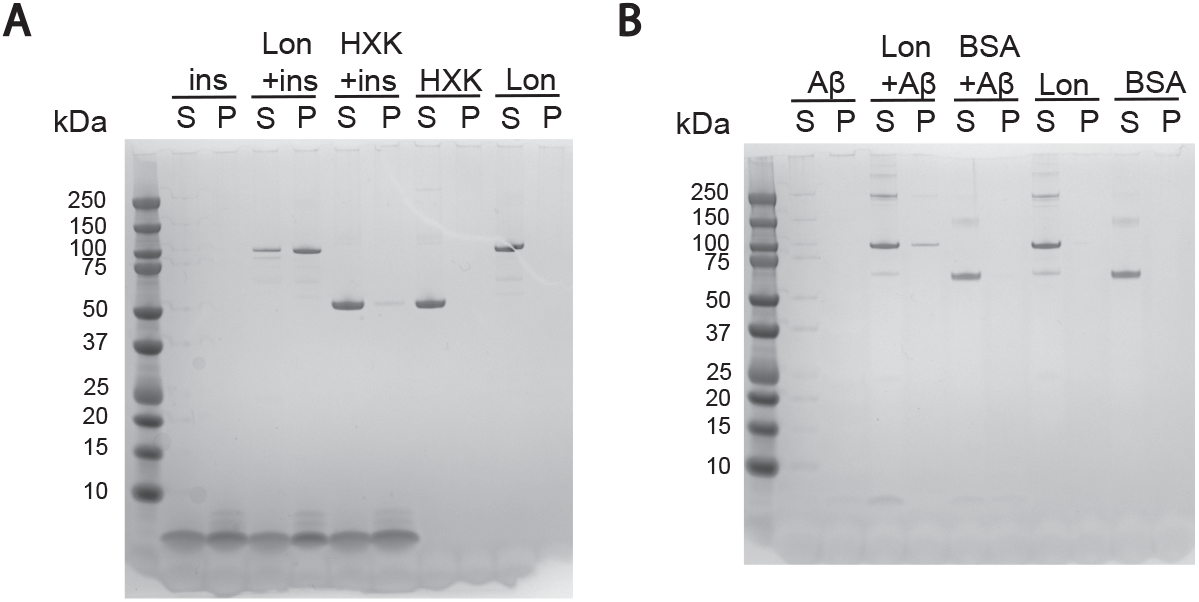
**A:** Finished fibrilization reactions of insulin (100 µM) in the presence of either hexokinase (1 µM, control) or proteolytically inactive LonP1 S855A (150 nM). **B:** Overnight incubation of 300 nM LonP1 or 600 nM BSA with 6 µM amyloid-beta 42. All reactions were separated in soluble (S) or precipitated (P) by centrigution at 13,000 × g for 10 minutes and analysed with SDS-PAGE.

**Supplementary figure S3:**
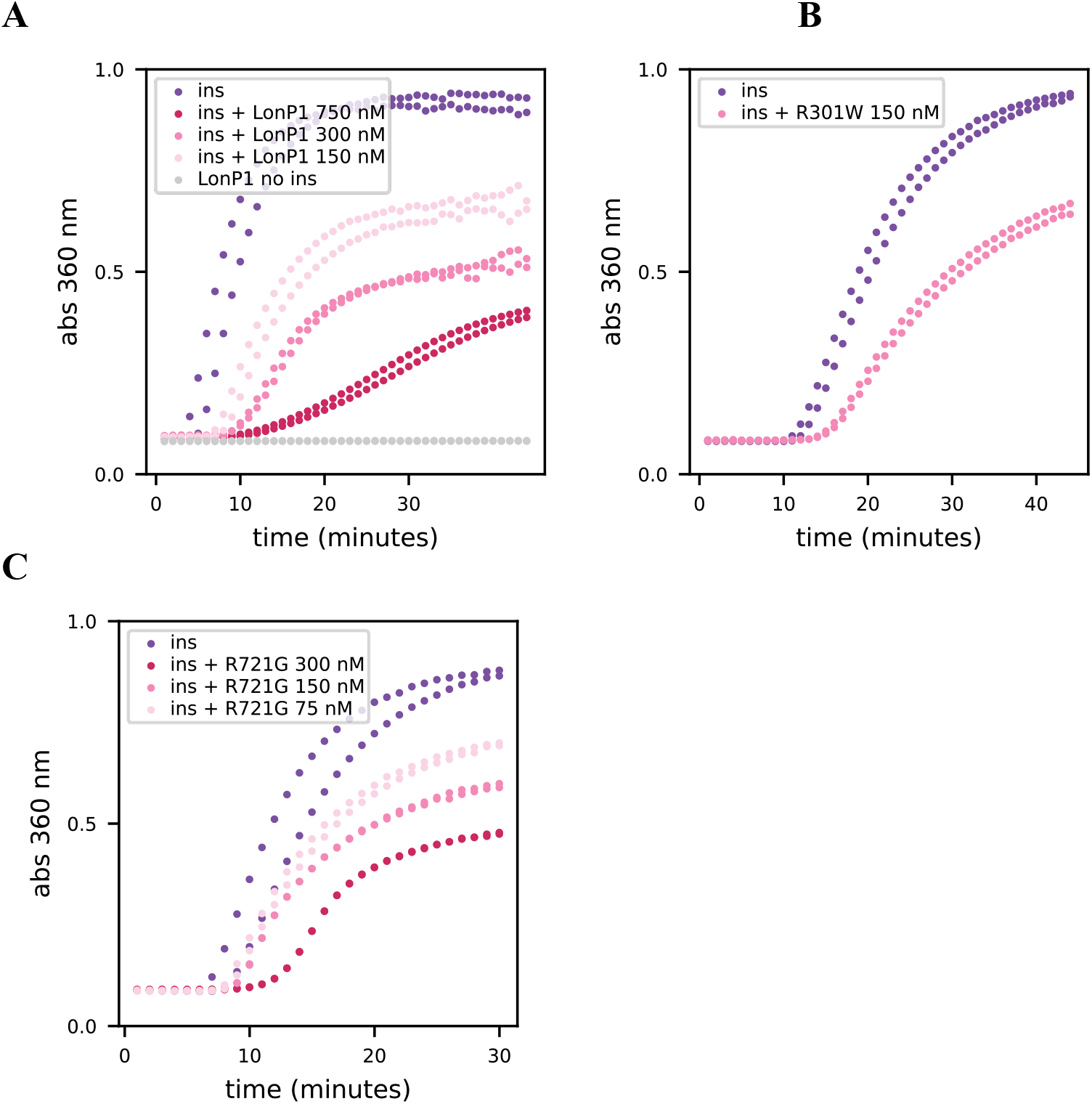
Insulin aggregation prevention of the LonP1 wild type (A) and variants R301W (B) and R721G (C). Assays were performed with LonP1 hexamers within one hour after Size Exclusion Chromotography.

**Supplementary figure S4:**
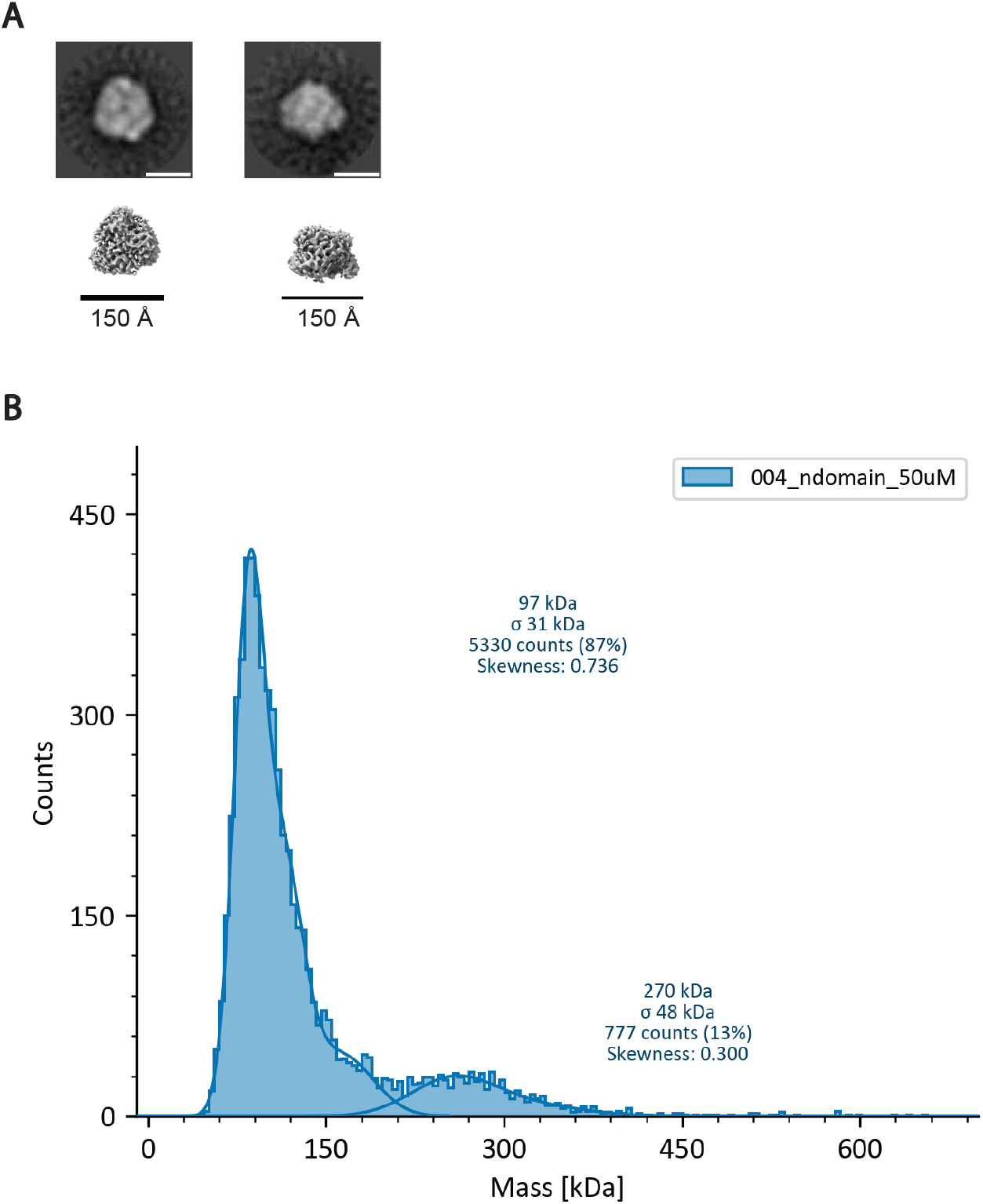
**A**. 2D-classes of negative stain electron microscopy of hexameric LonP1 N-domain particles, compared to a published electron microscopy map of the N-domain of the full-length LonP1 (pdb:7ngc), all scale bars are 150 Å. **B**. Mass photometry of the N-domain of LonP1 at oligomeric equilibrium, average of 3 measurements, at a sample concentration of 50 nM.

## References

1. N. Wang, S. Gottesman, M. C. Willingham, M. M. Gottesman, M. R. Maurizi, A human mitochondrial ATP-dependent protease that is highly homologous to bacterial Lon protease. Proc Natl Acad Sci U S A 90, 11247–11251 (1993).

2. C. K. Suzuki, K. Suda, N. Wang, G. Schatz, Requirement for the yeast gene LON in intramitochondrial proteolysis and maintenance of respiration. Science 264, 891 (1994).

3. L. Van Dyck, D. A. Pearce, F. Sherman, PIM1 encodes a mitochondrial ATP-dependent protease that is required for mitochondrial function in the yeast Saccharomyces cerevisiae. J Biol Chem 269, 238–242 (1994).

4. D. A. Bota, J. K. Ngo, K. J. Davies, Downregulation of the human Lon protease impairs mitochondrial structure and function and causes cell death. Free Radic Biol Med 38, 665–677 (2005).

5. G. K. Fu, D. M. Markovitz, The human LON protease binds to mitochondrial promoters in a single-stranded, site-specific, strand-specific manner. Biochemistry 37, 1905–1909 (1998).

6. Y. Matsushima et al., Mitochondrial Lon protease is a gatekeeper for proteins newly imported into the matrix. Commun Biol 4, 974 (2021).

7. C. S. Shin et al., LONP1 and mtHSP70 cooperate to promote mitochondrial protein folding. Nat Commun 12, 265 (2021).

8. M. Rep et al., Promotion of mitochondrial membrane complex assembly by a proteolytically inactive yeast Lon. Science 274, 103–106 (1996).

9. A. S. Olagunju et al., Mitochondrial dysfunction: A notable contributor to the progression of Alzheimer’s and Parkinson’s disease. Heliyon 9 (2023).

10. J. Lautenschläger et al., Intramitochondrial proteostasis is directly coupled to α-synuclein and amyloid β1-42 pathologies. J Biol Chem 295, 10138–10152 (2020).

11. W. Wang et al., Intraneuronal β-amyloid impaired mitochondrial proteostasis through the impact on LONP1. Proceedings of the Na@onal Academy of Sciences 120, e2316823120 (2023).

12. I. Mohammed et al., Catalytic cycling of human mitochondrial Lon protease. Structure 30, 1254–1268 e1257 (2022).

13. M. Shin et al., Structures of the human LONP1 protease reveal regulatory steps involved in protease activation. Nat Commun 12, 3239 (2021).

14. K. A. Strauss et al., CODAS syndrome is associated with mutations of LONP1, encoding mitochondrial AAA+ Lon protease. Am J Hum Genet 96, 121–135 (2015).

15. A. Besse, D. Brezavar, J. Hanson, A. Larson, P. E. Bonnen, LONP1 de novo dominant mutation causes mitochondrial encephalopathy with loss of LONP1 chaperone activity and excessive LONP1 proteolytic activity. Mitochondrion 51, 68–78 (2020).

16. Y. Matsushima, Y. Goto, L. S. Kaguni, Mitochondrial Lon protease regulates mitochondrial DNA copy number and transcription by selective degradation of mitochondrial transcription factor A (TFAM). Proc Natl Acad Sci U S A 107, 18410– 18415 (2010).

17. M. Falkenberg et al., Mitochondrial transcription factors B1 and B2 activate transcription of human mtDNA. Nat Genet 31, 289–294 (2002).

18. K. Pollecker, M. Sylvester, W. Voos, Proteomic analysis demonstrates the role of the quality control protease LONP1 in mitochondrial protein aggregation. J Biol Chem 297, 101134 (2021).

19. S. Kereiche et al., The N-terminal domain plays a crucial role in the structure of a full-length human mitochondrial Lon protease. Sci Rep 6, 33631 (2016).

20. J. Lee et al., Inhibition of mitochondrial LonP1 protease by allosteric blockade of ATP binding and hydrolysis via CDDO and its derivatives. Journal of Biological Chemistry 298, 101719 (2022).

21. G. Meisl et al., Molecular mechanisms of protein aggregation from global fikng of kinetic models. Nature Protocols 11, 252–272 (2016).

22. H. Stahlberg et al., Mitochondrial Lon of <i>Saccharomyces cerevisiae</i> is a ring-shaped protease with seven flexible subunits. Proceedings of the National Academy of Sciences 96, 6787–6790 (1999).

23. A. Y. Lee, C. H. Hsu, S. H. Wu, Functional domains of Brevibacillus thermoruber lon protease for oligomerization and DNA binding: role of N-terminal and sensor and substrate discrimination domains. J Biol Chem 279, 34903–34912 (2004).

24. M. R. Sawaya, M. P. Hughes, J. A. Rodriguez, R. Riek, D. S. Eisenberg, The expanding amyloid family: Structure, stability, function, and pathogenesis. Cell 184, 4857–4873 (2021).

25. R. Wasim, Bioenergetic failure and oxidative stress: mitochondrial contributions to Alzheimer’s disease. Inflammopharmacology 33, 5273–5289 (2025).

26. D. A. Bota, H. Van Remmen, K. J. A. Davies, Modulation of Lon protease activity and aconitase turnover during aging and oxidative stress. FEBS LeGers 532, 103–106 (2002).

27. R. Aishwarya et al., Diastolic dysfunction in Alzheimer’s disease model mice is associated with Aβ-amyloid aggregate formation and mitochondrial dysfunction. Scientific Reports 14, 16715 (2024).

28. A. Bezawork-Geleta, E. J. Brodie, D. A. Dougan, K. N. Truscott, LON is the master protease that protects against protein aggregation in human mitochondria through direct degradation of misfolded proteins. Sci Rep 5, 17397 (2015).

29. C. Chen et al., Parkinson’s disease neurons exhibit alterations in mitochondrial quality control proteins. npj Parkinson’s Disease 9, 120 (2023).

30. R. Shetty, R. Noland, G. Nandi, C. K. Suzuki, Powering down the mitochondrial LonP1 protease: a novel strategy for anticancer therapeutics. Expert Opin Ther Targets 28, 9–15 (2024).

31. Z. T. Farahbakhsh et al., Interaction of alpha-crystallin with spin-labeled peptides. Biochemistry 34, 509–516 (1995).

